# In-depth characterization of the cisplatin mutational signature in human cell lines and in esophageal and liver tumors

**DOI:** 10.1101/189233

**Authors:** Arnoud Boot, Mi Ni Huang, Alvin W.T. Ng, Szu-Chi Ho, Jing Quan Lim, Yoshiiku Kawakami, Kazuaki Chayama, Bin Tean Teh, Hidewaki Nakagawa, Steven G. Rozen

## Abstract

**Background and aims:** Cisplatin reacts with DNA, and thereby likely generates a characteristic pattern of somatic mutations, called a mutational signature. Despite widespread use of cisplatin in cancer treatment and its role in contributing to secondary malignancies, its mutational signature has not been delineated. We hypothesize that cisplatin’s mutational signature can serve as a biomarker to identify cisplatin mutagenesis in suspected secondary malignancies.
Knowledge of which tissues are at risk of developing cisplatin-induced secondary malignancies could lead to guidelines for non-invasive monitoring for secondary malignancies after cisplatin chemotherapy.

**Methods:** We performed whole genome sequencing of 10 independent clones of cisplatin-exposed MCF-10A and HepG2 cells, and delineated the patterns of single- and dinucleotide mutations in terms of flanking sequence, transcription strand bias, and other characteristics. We used the mSigAct signature presence test and non-negative matrix factorization to search for cisplatin mutagenesis in hepatocellular carcinomas and esophageal adenocarcinomas.

**Results:** All clones showed highly consistent patterns of single- and dinucleotide substitutions. The proportion of dinucleotide substitutions was high: 8.1% of single nucleotide substitutions were part of dinucleotide substitutions, presumably due to cisplatin’s propensity to form intra-and inter-strand crosslinks between purine bases in DNA. We identified likely cisplatin exposure in 9 hepatocellular carcinomas and 3 esophageal adenocarcinomas. All hepatocellular carcinomas for which clinical data were available and all esophageal cancers indeed had histories of cisplatin treatment.

**Conclusions:** We experimentally delineated the single- and dinucleotide mutational signature of cisplatin. This signature enabled us to detect previous cisplatin exposure in human hepatocellular carcinomas and esophageal adenocarcinomas with high confidence.

## Introduction

For 40 years, cisplatin and its derivatives have been cornerstones of the treatment of almost every type of cancer (Dasari and Tchounwou 2014; Dugbartey et al. 2016). However, cisplatin treatment often causes numerous side effects, including hepatotoxicity (Waseem et al. 2015; Dugbartey et al. 2016), and it increases the risk of developing secondary malignancies. For example, cisplatin based treatments almost always cure testicular cancers, but increase the risk of developing a solid tumor later in life 1.8-fold (Travis et al. 2005), and cisplatin treatment of several types of cancers increases the incidence of secondary leukemia’s (Ratain et al. 1987; Kushner et al. 1998). Cisplatin’s therapeutic properties depend partly on its DNA damaging activity, and the risk of secondary malignancies presumably stems from the consequent mutagenesis (Choi et al. 2014). This highlights the importance of understanding cisplatin mutagenesis and how it promotes carcinogenesis. This also highlights the need for a biomarker to identify cisplatin-induced secondary malignancies.

The mechanisms of cisplatin induced DNA damage have been extensively studied. When cisplatin enters the cells, its two chloride atoms are hydrolyzed, resulting in two positive charges (Masters and Koberle 2003; Behmand et al. 2015). Although the hydrolyzed molecule presumably reacts with many molecules in the cell, its therapeutic cytotoxicity is generally considered to stem from reactions with the N7 atoms of purine bases in DNA (Harrington et al. 2010; Dasari and Tchounwou 2014; Behmand et al. 2015). Most cisplatin-DNA adducts are crosslinks between two adjacent guanines (GpG, 65%) or between an adenine and a guanine (5’-ApG-3’, 25%). Mono-adducts and interstrand crosslinks are much rarer (Jamieson and Lippard 1999; Masters and Koberle 2003; Enoiu et al. 2012). Cisplatin induced DNA intrastrand crosslinks and mono-adducts are repaired through nucleotide excision repair (NER) (Zamble et al. 1996; Reardon et al. 1999; Hu et al. 2016). Interstrand crosslinks are the most difficult to repair and the most cytotoxic, because they covalently link the two strands of the DNA helix and consequently block transcription and replication (Jamieson and Lippard 1999; Masters and Koberle 2003; Enoiu et al. 2012; Hashimoto et al. 2016; Roy and Scharer 2016). The mechanisms of interstrand-crosslink repair have not yet been fully elucidated but appear to be complicated (Hashimoto et al. 2016; Roy and Scharer 2016).

Cisplatin likely causes a characteristic pattern of somatic mutations, known as a mutational signature, along with possible additional features such as fewer mutations on the transcribed strands of genes (Alexandrov et al. 2013a). Currently 30 mutational signatures are widely recognized, and they have a variety of known, suspected or unknown causes (Alexandrov et al. 2013a; Alexandrov et al. 2013b; Wellcome Trust Sanger Institute 2016). Mutational signatures can serve as biomarkers for endogenous mutagenic processes and exogenous exposures that led to the development of tumors.

We hypothesize that cisplatin’s mutational signature can serve as a biomarker to identify cisplatin mutagenesis in suspected secondary malignancies. Knowledge of which tissues are at risk of developing cisplatin-induced secondary malignancies could lead to guidelines for non-invasive monitoring for secondary malignancies after cisplatin chemotherapy.

Two previous studies investigated the mutational signature of cisplatin, one in *Caenorhabditis elegans* and one in a chicken (*Gallus gallus*) B-cell cell line (Meier et al. 2014; Szikriszt et al. 2016). Although both studies reported mutational signatures with primarily C>A mutations, the single-nucleotide substitution (SNS) signatures were otherwise dissimilar: the *C. elegans* signature was dominated by C<u>C</u>A>C<u>A</u>A and C<u>C</u>T>C<u>A</u>T mutations, while the chicken signature was dominated by <u>C</u>C><u>A</u>C mutations. This lack of similarity may have been due to the different model systems used, to the low numbers of mutations in the *C. elegans* study, or to experimental differences between the studies. In any case, these studies failed to unequivocally elucidate the mutational signature of cisplatin.

Therefore, we studied cisplatin mutations in MCF-10A, a non-tumorigenic human breast epithelial cell line, and in HepG2, a human liver cancer cell line. Here we report the extensive characterization of the cisplatin signature obtained, as well as its discovery in hepatocellular carcinomas and esophageal adenocarcinomas in patients previously exposed to cisplatin.

## Results

### Cisplatin’s single-nucleotide substitution signature

We exposed two independent cultures of MCF-10A cells to 0.5 μM and 1 μM, and one culture of HepG2 to 0.75 μM of cisplatin once a week for 8 weeks. Single cells were isolated and expanded for whole-genome sequencing and mutational analysis. We sequenced the untreated cell lines, 3 MCF-10A clones for each concentration (one exposed for 4 weeks and 2 exposed for 8 weeks) and 4 HepG2 clones (exposed for 8 weeks). Mean coverage was >33x, and in total we identified 70,313 single nucleotide substitutions (SNSs) (Supplemental Table S1).

The SNS mutation spectra from all clones were highly similar (Figure 1A, Supplemental Fig. S1A, Supplemental Table S2, all Pearson correlations > 0.958 and cosine similarities > 0.971). The most prominent features were two C>T peaks (C<u>C</u>C>C<u>T</u>C and C<u>C</u>T>C<u>T</u>T) and four T>A peaks (C<u>T</u>>C<u>A</u>). There were also substantial numbers of C>A mutations (~26.0% of all mutations), and peaks at G<u>C</u>C>G<u>A</u>C and G<u>C</u>C>G<u>G</u>C. Figure 1B and Supplemental Fig. S1B display the signatures as mutation rates per trinucleotide, which better reflects the sequence specificity of mutational processes because they are not affected by differences in trinucleotide abundances. For example, Figure 1B shows more prominent C<u>C</u>C>C<u>T</u>C peaks and reveals that the gap at C<u>C</u>G>C<u>T</u>G in Figure 1A reflects the low abundance of CCG trinucleotides in the genome rather than reduced mutagenicity.

**Figure 1:**
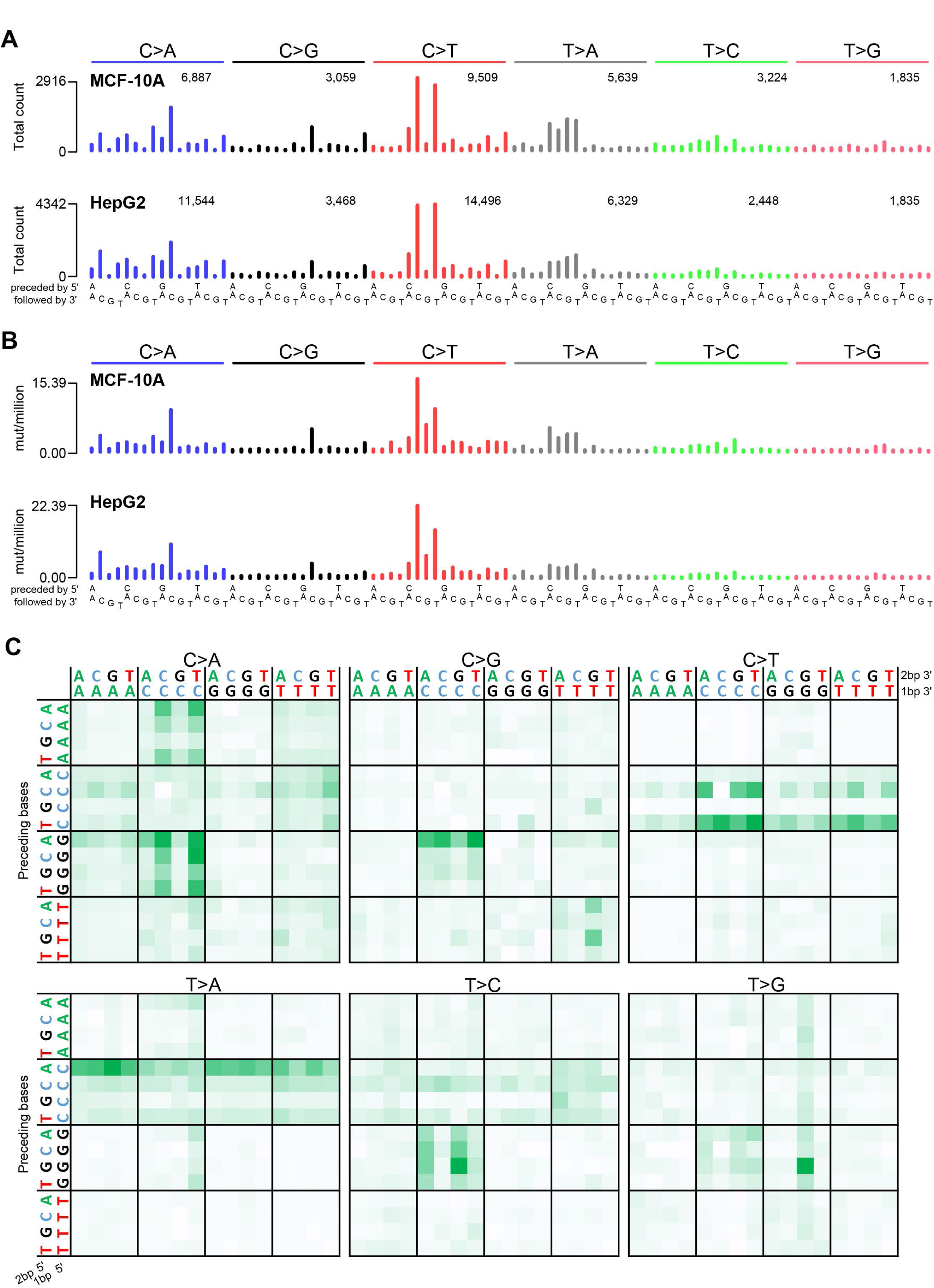
Cisplatin mutational signature. Trinucleotide-context mutational spectra shown as (**A**) raw counts and (**B**) rate of mutations per million trinucleotides for all MCF-10A (top panel) and all HepG2 (bottom panel) clones combined. In (**A**), the number of mutations per SNS type is shown above the corresponding bars. (**C**) Pentanucleotide sequence contexts for all samples combined, normalized by pentanucleotide occurrence in the genome. See also Supplemental Fig. S2.

In addition to consistent patterns of the bases immediately 5’ and 3’ of cisplatin SNSs, there were also many preferences 2 bp 5’ and 3’ of the SNSs (Figure 1C, Supplemental Fig. S2). For example, C<u>T</u>>C<u>A</u> mutations were usually preceded by an A (AC<u>T</u>>AC<u>A</u>). Similarly, C<u>C</u>>C<u>T</u> mutations were usually preceded by a pyrimidine (YC<u>C</u>>YC<u>T</u>). These and other preferences at the -2 bp or +2 bp positions were statistically significant (Supplemental Fig. S3). Examination of the -3 bp or +3 bp positions of SNSs revealed no additional sequence context preferences (Supplemental Fig. S4).

### Associations of cisplatin-induced single-nucleotide substitutions with genomic features

Many mutational processes cause fewer mutations due to damage on the transcribed strands of genes than on the non-transcribed strands. This is termed transcription strand bias and is due to transcription-coupled nucleotide excision repair (TC-NER) of adducted bases in the transcribed (antisense) strands. Since cisplatin forms adducts on purines, we would expect reduced numbers of mutations when G and A is on the transcribed strand (corresponding to C and T on the sense strand). As expected, C>A, C>T and T>A SBSs were strongly reduced on the sense strand (Supplemental Fig. S5) (Fousteri and Mullenders 2008; Harrington et al. 2010; Dasari and Tchounwou 2014; Behmand et al. 2015; Hu et al. 2016). Also consistent with TC-NER, strand bias for C>A, C>T and T>A mutations was stronger in more highly expressed genes (*p* = 1.45×10^-50^ and 1.20×10^-116^, one-sided Chisquared test for all MCF-10A and for all HepG2 clones combined, respectively, Figure 2A, Supplemental Fig. S6). Finally, TC-NER efficiency decreases from the 5’ to the 3’ ends of transcripts (Conaway and Conaway 1999; Hu et al. 2015; Huang et al. 2017). Consistent with this, strand bias for C>A, C>T and T>A SNSs decreased toward the 3’ ends of transcripts (*p* = 2.46×10^-12^ and *p* = 1.85×10^-12^, logistic regression for all MCF-10A clones and for all HepG2 clones combined, respectively, Figure 2B, Supplemental Fig. S7).

**Figure 2:**
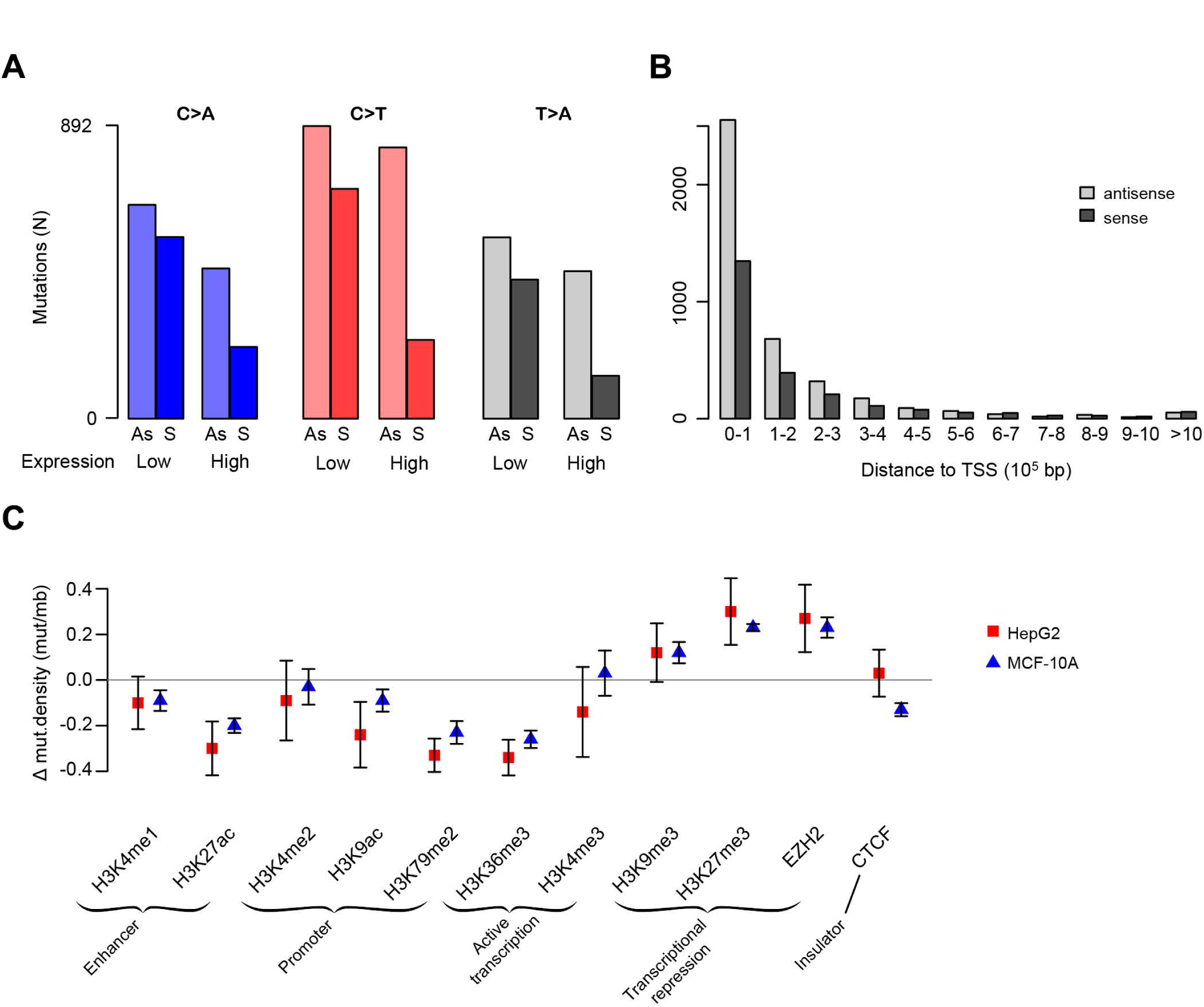
Associations between cisplatin mutagenesis intensity and genomic features. (**A**) Transcription strand bias is more prominent in highly expressed genes for C>A, C>T and T>A mutations. See also Supplemental Fig. S6. (**B**) Transcription strand bias decreases with increasing distance from the transcription start site (TSS). See also Supplemental Fig. S7. Mutations were binned per 100,000bp, i.e. the first bars are the numbers of mutations within the first 100,000bp from the TSS, the next bars are the numbers of mutations in the region from 100,001 to 200,000bp from the TSS, and so on. (**C**) Mutation density in regions with histone modifications and in binding sites for EZH2 and CTCF. The y axis is the mean mutation density for the given region relative to the mutation density of each respective sample; bars show standard error of the mean (Supplemental Table S1).

In addition to transcription strand bias, cisplatin mutagenesis also showed replication timing bias, with a higher mutation density in late replicating regions (*p* = 9.39×10^-73^ and 1.96×10^-136^ binomial test for MCF-10A and HepG2 clones, respectively). We noted high variability in replication timing bias between the different clones, the cause of which remains unclear. Interestingly, C>T mutations showed lower replication timing bias than other mutations classes (Supplemental Fig. S8). There was no difference in mutation density between leading- and lagging replication strands.

For some mutational processes, mutagenesis intensity varies by chromatin state (Polak et al. 2015; Seplyarskiy et al. 2015; Kaiser et al. 2016). Additionally, there is increased cisplatin adduct formation in open chromatin compared to closed chromatin (Hu et al. 2016). In both cell lines, regions containing active promoters, enhancers and actively transcribed genes were less highly mutated, and regions associated with heterochromatin and transcriptional repression were more highly mutated (Figure 2C).

### Cisplatin’s dinucleotide substitution signature

To investigate the presence of dinucleotide substitutions (DNSs) in the cisplatin genomes we selected all adjacent SNS, and verified that both SNS were on the same reads (see Materials and Methods). We identified 2,839 DNSs in the cisplatin genomes, of which most were mutations from CC, CT, TC and TG (Figure 3A, Supplemental Fig. S9). We hypothesized that mutations from CC, CT, and TC are consequences of intrastrand crosslinks at GpG, ApG and GpA, and that mutations from TG were consequences of diagonally-offset interstrand guanine-adenine crosslinks 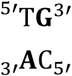 (crosslinked bases in bold). Mutations from AT, TA and TT were rare, which is consistent with previous reports that cisplatin does not induce adenine-adenine crosslinks (Supplemental Table S3) (Jamieson and Lippard 1999; Masters and Koberle 2003).

**Figure 3:**
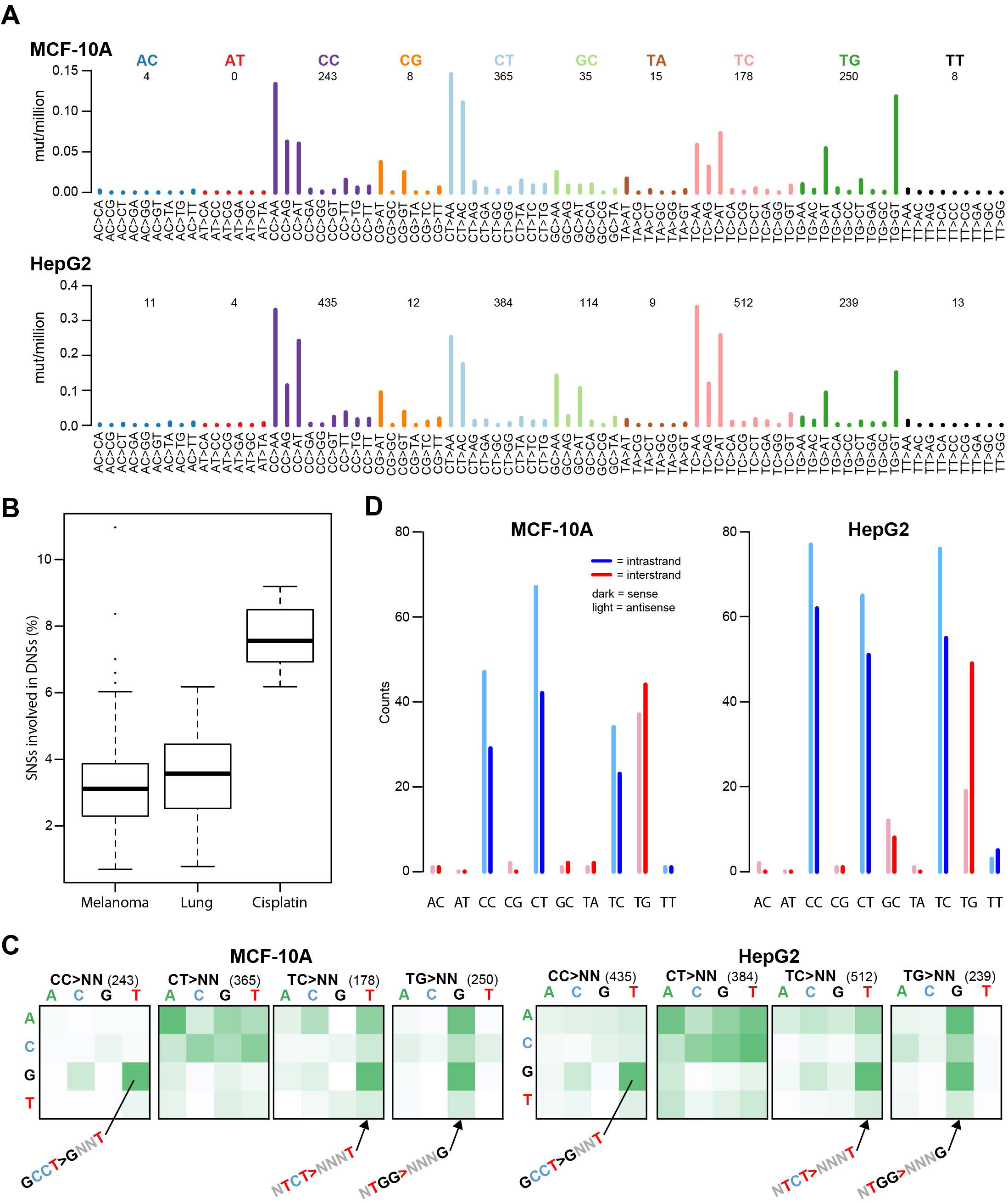
Cisplatin induced dinucleotide substitutions (DNSs). (**A**) DNS mutation spectra of all MCF-10A (top panel) and all HepG2 (bottom panel) clones combined, displayed as DNSs per million dinucleotides (i.e. normalized for dinucleotide abundance in the genome). (**B**) Cisplatin induces higher numbers of DNSs than other mutational processes associated with dinucleotide substitutions such as UV (melanoma) and smoking (lung). (**C**) ±1bp sequence context preferences for the most prominent DNS mutation classes (CC>NN, CT>NN, TC>NN and TG>NN). The total number of DNSs per mutation class is indicated in parentheses. The vertical axis is the preceding (5’) base, the horizontal axis is the following (3’) base. Some prominent enrichments in sequence context are indicated (G<u>CC</u>T>G<u>NN</u>T, N<u>TC</u>T>N<u>NN</u>T and N<u>TG</u>G>N<u>NN</u>G). The full sequence context preference plots, both raw counts and normalized for tetranucleotide abundance in the genome are shown in Supplemental Fig. S10. (**D**) Transcription strand bias of dinucleotide substitutions. Potential intrastrand crosslink sites are shown in blue, potential interstrand crosslink sites are shown in red.

The proportion of SNSs involved in DNSs ranged from 6.2% to 9.2%. To relate this to other mutagenic processes known to be associated with DNSs, we examined the percentage of SNSs involved in DNSs associated with COSMIC Signatures 4 (smoking-related) and 7 (due to UV exposure) (Wellcome Trust Sanger Institute 2016). We studied Signature 4 in 24 lung adenocarcinomas (Imielinski et al. 2012) and Signature 7 in 112 melanomas (Zhang et al. 2011). In both tumor types, the percentage of SNSs involved in DNSs was significantly lower than in cisplatin (Figure 3B, mean 3.5%, sd=1.4%, *p*=6.5×10^-10^ and mean=3.3%, sd=1.6%, *p*=4.6×10^-14^ respectively, 2-sided t-tests versus cisplatin). We hypothesize that this high proportion of DNSs in cisplatin stems from cisplatin’s propensity to form intrastrand crosslinks between adjacent bases and to form diagonally offset interstrand crosslinks.

To investigate possible sequence context preferences of cisplatin DNSs, we plotted 1bp contexts of each reference dinucleotide, irrespective of the mutant allele (Figure 3C, Supplemental Fig. S10). There was strong enrichment for TC and TG DNSs in <u>TC</u>T and <u>TG</u>G contexts. Both TC and TG DNSs were further enriched for a 5’ flanking purine (Supplemental Fig. S10, S11). The strongest sequence context preference was for CC>NN mutations, 49.8% of which occurred in the G<u>CC</u>T context (Supplemental Fig. S10, S11). As methodological controls, we also evaluated ±1bp sequence context for DNSs associated with COSMIC Signatures 4 and 7. DNSs associated with Signature 7 showed strong sequence context preference for most mutation classes, including CC>NN, CT>NN and TT>NN (Supplemental Fig. S12). The context preferences were very different however, from those of cisplatin DNSs. By contrast, DNSs associated with Signature 4 had only weak sequence context preferences (Supplemental Fig. S12).

### Associations of cisplatin-induced dinucleotide substitutions with genomic features

To assess transcription strand bias in DNSs, we examined separately the mutations hypothetically involving interstrand purine-purine crosslinks, predominantly mutations from the 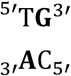 configuration, and the mutations hypothetically involving intrastrand purine-purine crosslinks (predominantly mutations from CC, CT, and TC). We observed transcription strand bias at the potential intrastrand crosslink sites other than TC in most of the MCF-10A and HepG2 clones. (Figure 3D, Supplemental Fig. S13). There was no consistent evidence of transcription strand bias at potential interstrand crosslink sites (mainly TG) in the MCF-10A clones. However, for 3 of the 4 HepG2 clones, there were fewer mutations when TG was on the transcribed (antisense) strand (Supplemental Fig. S13). As methodological controls, we also evaluated transcription strand bias for DNSs associated with COSMIC Signatures 4 and 7, in which we also detected strand bias (Supplemental Fig. S14).

With respect to other genomic features, the replication-timing bias of DNSs was similar to that of the SNSs (Supplemental Fig. S8). Association of DNS density with marks of active and repressed chromatin was similar to that of SNS density, with the following exceptions (Supplemental Fig. S15). DNS density was markedly higher than SNS density in regions of H3K9 acetylation and markedly lower at binding sites of EZH2 (enhancer of zeste homologue 2 polycomb repressive complex 2 subunit). In addition, DNS density was markedly higher than SNS density at binding sites of CTCF (CCCTC-binding factor).

### Other mutation types

We also examined small insertion and deletion mutations (indels), copy number alterations, and structural variants in the cisplatin exposed MCF-10A and HepG2 clones. We identified 4,208 indels in the cisplatin exposed clones. The indels were unremarkable, consisting primarily of single-nucleotide insertions or deletions (~78%, Supplemental Fig. S16). Like SNSs, indels were enriched in late replicating regions (Supplemental Fig. S8). The distribution of indels with respect to other genomic features was very similar to that of DNSs (Supplemental Fig. S15). There were very few copy number alterations or structural variants (Supplemental Fig. S17, S18), suggesting that cisplatin did not induce detectable genomic instability.

### Likely cisplatin mutational signature in human tumors

We examined publicly available human tumor mutation data for evidence of the experimental cisplatin signature. Notably, mutational signature W6, which was reported in the whole genome sequences of hepatocellular carcinomas (HCCs), resembles the experimental cisplatin signature (cosine similarity = 0.781, Supplemental Fig. S19) (Fujimoto et al. 2016). Although the relative proportions of the major substitution classes (C>A, C>T and T>A) are rather different between Signature W6 and our experimental cisplatin signature, the profiles within each mutation class are similar (cosine similarities for C>A, C>T and T>A of 0.915, 0.917 and 0.981, respectively, Supplemental Fig. S19). Given this resemblance, we searched for the cisplatin SNS signature using the mSigAct signature presence test (see Materials & Methods (Ng et al. 2017)) in data from Japanese and Hong Kong HCCs (Kan et al. 2013; Fujimoto et al. 2016). Out of 342 HCCs, 10 showed evidence of cisplatin exposure (Table 1, Figure 4A, Supplemental Fig. S20, compare with Figure 1A). To further assess presence of cisplatin mutagenesis, we also examined the dinucleotide spectra of these samples (Figure 4B, Supplemental Fig. S21, compare with Figure 3A). 7 of the 10 HCCs with the cisplatin SNS spectrum also had high cosine similarities between their DNS spectra and the cisplatin signature (Figure 4C) and high numbers of DNSs relative to their total SNS load (ranging from 2.9 to 6.2%, compared to the median of 1.6%, for all HCCs, Supplemental Fig. S22A).

**Table 1:** HCCs and ESADs with cisplatin‐associated mutagenesis

| Tumor ID | Cancer type | Total SNSs | Cisplatin SNSs | mSigAct P* | DNSs | DNS cosine similarity to experimental signature | DNSs due to cisplatin** | Conclusion based on SNS and DNS analysis | Patient history*** | Reference |
| --- | --- | --- | --- | --- | --- | --- | --- | --- | --- | --- |
| HK034 | HCC | 7844 | 2274 | 1.3E-11 | 153 | 0.827 | 130 | Cisplatin positive | NA | Kan et al., 2013 |
| RK028 | HCC | 20792 | 7594 | 1.5E-21 | 642 | 0.773 | 533 | Cisplatin positive | cisplatin DEB-TACE | Fujimoto et al., 2016 |
| RK047 | HCC | 8345 | 1522 | 1.1E-04 | 73 | 0.498 | 8 | Negative | No neo-adjuvant chemotherapy | Fujimoto et al., 2016 |
| RK056 | HCC | 17085 | 7097 | 4.4E-26 | 479 | 0.874 | 426 | Cisplatin positive | cisplatin DEB-TACE | Fujimoto et al., 2016 |
| RK072 | HCC | 8893 | 1130 | 4.9E-04 | 92 | 0.698 | 50 | Cisplatin positive | cisplatin DEB-TACE (5 rounds) | Fujimoto et al., 2016 |
| RK074 | HCC | 22406 | 6903 | 2.5E-18 | 476 | 0.865 | 415 | Cisplatin positive | cisplatin DEB-TACE | Fujimoto et al., 2016 |
| RK140 | HCC | 10132 | 1115 | 2.5E-04 | 125 | 0.800 | 73 | Cisplatin positive | cisplatin DEB-TACE, 4 years, 1 year and 6 months prior | Fujimoto et al., 2016 |
| RK205 | HCC | 10406 | 1751 | 3.0E-06 | 158 | 0.720 | 150 | Cisplatin positive | cisplatin DEB-TACE, prior history of HCC, resected 27 months ago | Fujimoto et al., 2016 |
| RK223 | HCC | 10680 | 1157 | 1.4E-04 | 141 | 0.640 | 65 | Negative | No neo-adjuvant chemotherapy | Fujimoto et al., 2016 |
| RK241 | HCC | 10610 | 3373 | 5.1E-16 | 235 | 0.767 | 177 | Cisplatin positive | cisplatin DEB-TACE, prior history of colorectal cancer | Fujimoto et al., 2016 |
| RK256 | HCC | 11240 | 2530 | 2.4E-09 | 167 | 0.806 | 142 | Cisplatin positive | cisplatin DEB-TACE, prior history of HCC, resected 37 and 18 months ago | Fujimoto et al., 2016 |
| RK309 | HCC | 4785 | 1379 | 2.3E-05 | 12 | 0.558 | To few DNSs to analyze | Negative | No neo-adjuvant chemotherapy | Fujimoto et al., 2016 |
| SA594320 | ESAD | 23483 | 5180 | 9.6E-19 | 313 | 0.902 | 266 | Cisplatin positive | Cisplatin treated | Noorani et al., 2017 |
| SA594557 | ESAD | 7433 | 763 | 1.0E-05 | 63 | 0.877 | 41 | Cisplatin positive | Cisplatin treated | Noorani et al., 2017 |
| SA594775 | ESAD | 14967 | 1914 | 4.3E-06 | 124 | 0.790 | 58 | Cisplatin positive | Cisplatin treated | Noorani et al., 2017 |

\* bonferroni level of significance is 1.5E-04 (0.05/342) for the HCCs and 3.6E-04 (0.05/140) for the ESADs
\*\* DNS assignment by ssNMF. The cisplatin DNS signature was given as input, and 2 other DNS signatures were requested. Reported here is the number of DNSs assigned to the cisplatin DNS signature.
\*\*\* NA denotes data not available

**Figure 4:**
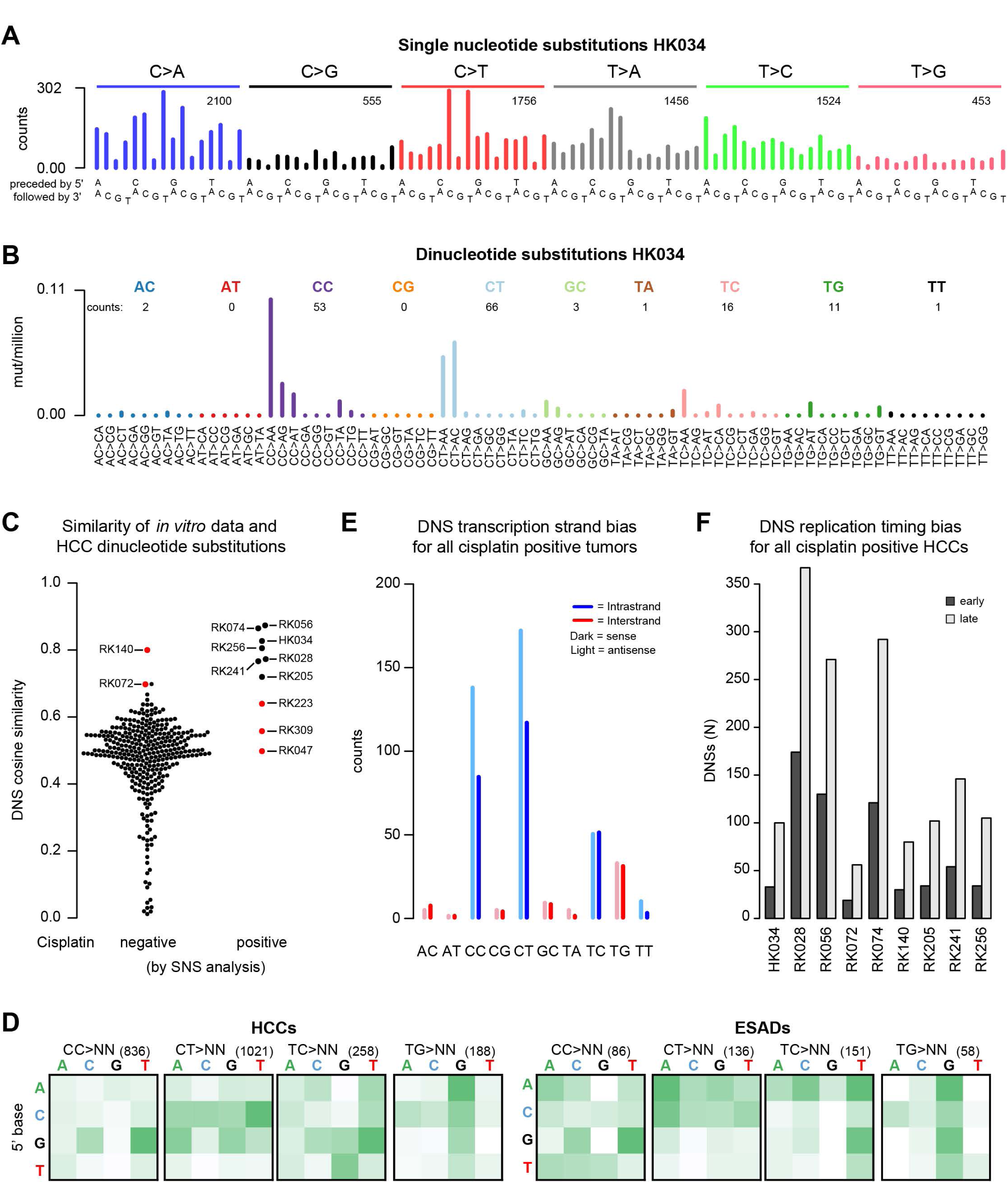
Cisplatin mutational signature in human hepatocellular carcinomas (HCCs) and esophageal adenocarcinomas (ESADs). (**A**) Example SNS and (**B**) DNS mutational spectra of a tumor that tested positive for the cisplatin signature in the SNS analysis (HK034). In (**A**) and (**B**), numbers of mutations in each mutation class are indicated. (**C**) DNS cosine similarities between the experimental cisplatin signature and HCCs, grouped on whether they were negative (left) or positive (right) for cisplatin mutagenesis in the SNS analysis. Red dots represent HCCs that were found positive for cisplatin mutagenesis in the SNS analysis but did not show the cisplatin DNS signature (false-positives) and samples that were not found cisplatin positive in the SNS analysis but were concluded to be cisplatin positive based on the DNS analysis (false-negatives). (**D**) ±1bp sequence context preferences for the most prominent DNS mutation classes in cisplatin-positive HCCs and ESADs. Total numbers of DNSs per mutation class are indicated in parentheses. The vertical axis is the preceding (5’) base, the horizontal axis is the following (3’) base. (**E**) DNS transcription strand bias in all cisplatin-positive tumors combined. For the individual sample plots, see Supplemental Fig. S27. (**F**) DNS replication timing bias in cisplatin-positive HCCs. DNSs were classified as being in either early or late replicating regions as described in Materials & Methods.

We also analyzed the mutational spectra of 140 esophageal adenocarcinomas (ESADs), of which 68 had been treated with cisplatin prior to surgery (Noorani et al. 2017). SNS analysis suggested that 3 of the cisplatin treated ESADs had the cisplatin signature, whereas we found no evidence of cisplatin mutagenesis in any of the untreated ESADs. The DNS analysis supported likely cisplatin exposure in all ESADs identified in the SNS analysis (Table 1, Supplemental Fig. S22B, S23, S24).

We further investigated whether DNS analysis could identify cisplatin-exposed tumors that were missed by the SNS analysis. We performed semi-supervised nonnegative matrix factorization (ssNMF) on all tumors with ≥25 DNSs, specifying the cisplatin DNS signature as one input signature and asking for discovery of 1 to 4 additional signatures (Materials & Methods, Supplemental Fig. S25, S26). All 7 previously identified cisplatin-positive HCCs had >50% DNS attributed to cisplatin by ssNMF, as did an additional 13 HCCs. Among these 13 HCCs, two, RK072 and RK140, had high cosine similarities with the experimental cisplatin DNS signature and had relatively high proportions of DNSs compared to SNSs (Table 1, Supplemental Fig. S22A). Although the SNS based *p* values were not significant after multiple-testing correction, we nevertheless concluded based on the combined SNS and DNS analyses that RK072 and RK140 showed strong evidence for cisplatin mutagenesis. For the remaining 11 HCCs with >50% cisplatin-associated DNSs, neither mSigAct nor visual inspection of the SNS spectra warranted reclassification as cisplatin positive.

Use of ssNMF also identified high proportions of cisplatin-associated DNSs in several ESADs. These included the 3 identified in our initial analysis. Of the remainder, neither the mSigAct signature presence test on the SNSs nor visual inspection of the SNS spectra warranted reclassification as cisplatin positive. None of the chemotherapy naïve ESADs displayed signs of cisplatin mutagenesis based on the DNS analysis.

Beyond the mutation frequency spectrum, the other characteristics of the DNSs in the cisplatin positive HCCs and ESADs were very similar to DNS characteristics in the experimental data. First, the DNS sequence context preference of the cisplatin-positive HCCs and ESADs was extremely similar to the experimental data (Figure 4D, compare with Figure 3C). The TC and TG DNSs, were less frequent in the tumors than in the experimental data, but nevertheless showed very similar sequence context preferences. Second, like the cisplatin exposed cells, most HCCs and ESADs showed strong transcription strand bias at CC and CT DNSs but not at TC DNSs (Figure 4E, Supplemental Fig. S27). Also like the cisplatin exposed MCF-10A clones, none of the HCCs and ESADs had detectable transcription strand bias at potential interstrand crosslink sites (mainly TG DNSs, Supplemental Fig. S27). Third, like the experimental data, the DNSs of the cisplatin positive tumors did not show replication strand bias but did show strong replication timing bias (Figure 4F).

The clinical records of the Japanese HCCs (Fujimoto et al. 2016) confirmed cisplatin exposure of all of the 7 HCCs identified positive for the cisplatin mutational signature (Table 1). All 7 had received cisplatin-based transarterial chemoembolization using drug eluting beads (DEB-TACE) several months prior to surgical resection. In addition to DEB-TACE treatment of the sampled tumor, patients RK205, RK241 and RK256 also had had prior malignancies (Table 1). The variant allele frequencies of the cisplatin-associated DNSs were similar to the variant allele frequencies of all SNSs, including those not likely due to cisplatin exposure (Supplemental Table S4). This suggested that the cisplatin was an early event in tumorigenesis, which would be concordant with rapid clonal expansion after DEB-TACE treatment (Zen et al. 2011). Notably, the 3 HCCs (RK047, RK223 and RK309) that we suspected to be false-positives based on DNS analysis had no record of treatment with cisplatin prior to surgery.

## Discussion

We have delineated the *in vitro* multidimensional mutational signature of cisplatin in two human cell lines. This comprised extensive characterization of patterns of SNSs in tri-and pentanucleotide contexts and the associations of SNSs with genomic features. We also found patterns of DNSs and flanking bases and the associations of DNSs with genomic features that were highly informative. We began with *in vitro* delineation because it directly links mutational signatures to etiologies and because it generates signatures that are relatively unobscured by other mutational processes. We analyzed whole genome data because these provide >50 times more mutations than exomes and consequently greater stability and reproducibility of signatures. Indeed, whole genome data are practically essential for analysis of DNSs, which are rare compared to SNSs. Importantly, with the experimentally delineated SNS and DNS signatures in hand, we were able to detect cisplatin mutagenesis in HCCs and ESADs with high confidence. All HCCs for which clinical data were available and all esophageal cancers indeed had histories of prior cisplatin treatment. We therefore conclude that the mutational signature established here serves as a biomarker for cisplatin mutagenesis that could be used to determine whether or not a suspected secondary malignancy was indeed induced by cisplatin.

Prior to this study, 2 different experimentally elucidated mutational signatures of cisplatin were reported, one in *Caenorhabditis elegans,* and the other in cultured chicken B-cells (DT40) (Meier et al. 2014; Szikriszt et al. 2016). Both studies found primarily C>A mutations, but in terms of SNSs in trinucleotide context, the signatures bore no resemblance to each other or to the MCF-10A/HepG2 signature reported here (Supplemental Fig. S28A). In the *C. elegans* data, this was true for both the DNA repair proficient worms as well as for all worms combined. Like our experimental data, the exposed worms and DT40 cells had relatively high numbers of DNSs relative to SNSs, and strikingly, the DT40 DNS spectra closely resembled our experimental DNS signature (cosine similarity = 0.935, Supplemental Fig. S28B). However, in neither system was it possible to discern the MCF-10A/HepG2 SNS signature in the mutation spectra, due to the high number of C>A mutations (Supplemental Fig. S28A). We also note that the C>A mutations in the treated worms and DT40 cells do not resemble any currently known mutational signature or artefact (Wellcome Trust Sanger Institute 2016). In light of the similarity between the MCF-10A/HepG2 and DT40 DNS signatures, we further investigated whether the DT40 SNS signature might be present in HCCs or ESADs. Comparisons using the mSigAct signature presence test concurred that compared to the DT40 SNS signature, the MCF-10A/HepG2 signature is more effective at detecting cisplatin-mutagenized HCCs and ESADs and at explaining their mutational spectra (Supplemental Data S1).

The differences between the MCF-10A/HepG2 SNS cisplatin signature and the *C. elegans* and DT40 signatures might stem from the different model organisms used, which may differ in DNA damage susceptibility and characteristics of DNA repair and replication errors. In any case, the differences between the previously published cisplatin spectra and the MCF-10A/HepG2 signature emphasize the need for standardization of *in vitro* mutational signature models. We propose that it is prudent to use human cell lines for experimental elucidation of mutational signature etiology, to avoid possible differences in translesion synthesis and DNA repair proficiencies between organisms.

Mutational processes reflect the cumulative effect of 3 steps: (i) DNA damage (for cisplatin, adduct formation), (ii) DNA repair (for cisplatin, NER), which may or may not correct the damage, and (iii) if DNA repair fails, translesion synthesis across the damaged base or bases, which may replicate the DNA correctly or incorrectly, in the latter instance creating a mutation. In this study, while known patterns of adduct formation did not predict the patterns of substitutions (Figure 5), we can nevertheless postulate models that explain the observed mutations by combining our knowledge of adduct formation and models of how DNA replication and translesion synthesis might behave (II and III).

**Figure 5:**
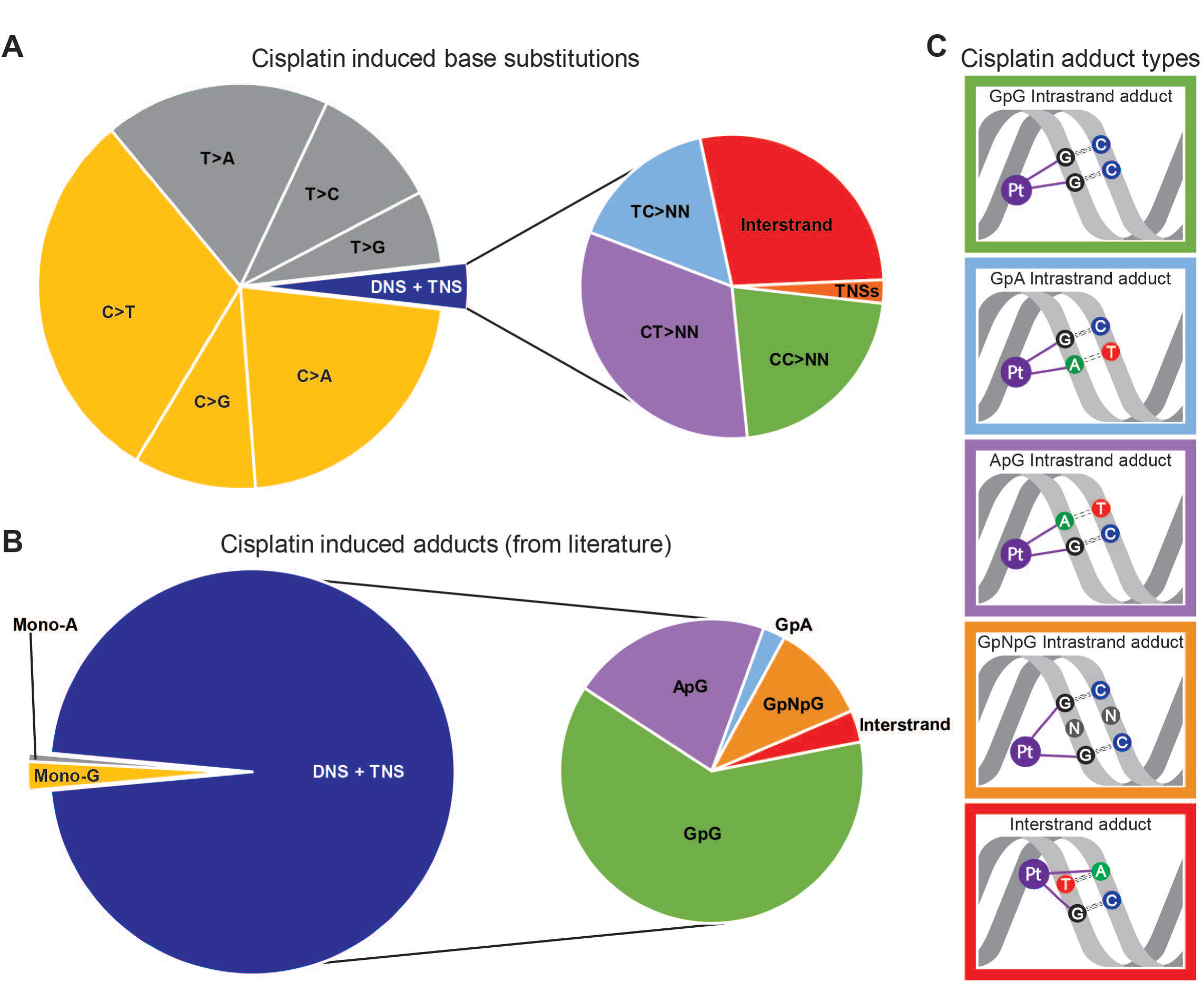
Comparison of proportions of cisplatin-induced substitutions and reported cisplatin adducts. (**A**) Relative abundance of cisplatin-induced base substitutions in the experimental signature. TNS = trinucleotide substitutions. (**B**) Relative abundances of cisplatin-adducts from: (Eastman 1983; Fichtinger-Schepman et al. 1989; Jamieson and Lippard 1999; Baik et al. 2003; Enoiu et al. 2012). Colors of mutations in **A** correspond to colors of the adducts they are expected to be caused by in **B**. (**C**) Schematic representations of adducts in B related to cisplatin-induced substitutions in **A**: The colors of the borders of the schematic adduct representation correspond to the colors used in the zoomed-in section of the pie-charts on the right sides of **A** and **B**

First, despite high proportions of DNSs relative to SNSs, SNSs still greatly outnumbered the DNSs (Figure 5A). We postulate that these SNSs are formed by correct translesion synthesis opposite one of the purines of the purine-purine intrastrand crosslinks, and misincorporation occurring opposite the other, as has been shown for UV-induced intrastrand crosslinks (McCulloch et al. 2004). This is supported by the high number of SNSs at potential intrastrand crosslink sites: 85% of the SNSs are at GpG, GpA or ApG sites (Supplemental Fig. S29). Closer inspection of SNSs in trinucleotides encompassing only a single potential intrastrand crosslink site revealed that at most such sites, SNSs are more common at the 3’ adducted-base across every cell-line clone (Supplemental Fig. S30). However, at potential adenine-guanine intrastrand crosslink sites, SNSs are more common at the 5’ base. We do not have an explanation for this difference. Possibly different translesion synthesis polymerases are involved in traversing the various intrastrand crosslinks.

Second, the relative abundance of the different types of DNSs did not correspond to the reported ratios of intrastrand and interstrand adducts at their respective dinucleotides (compare the right pie-charts of Figures 5A,B, with graphical representations of the most prominent adducts in Figure 5C). 24.7% of DNSs were in potential interstrand crosslink sites, while these represent <5% of cisplatin-adducts (Jamieson and Lippard 1999; Enoiu et al. 2012). However, the higher proportion of DNSs putatively due to interstrand crosslinks is consistent with interstrand crosslinks being more damaging and harder to repair than intrastrand crosslinks (Andreassen and Ren 2009; Hashimoto et al. 2016; Roy and Scharer 2016).

In this study, combined SNS and DNS information was crucial for high-confidence detection of cisplatin mutagenesis in human tumors. SNS analysis alone would have identified 3 false-positives and missed RK072 and RK140, and DNSs analysis alone would have identified several likely false positives. Ideally, the field of mutational signature analysis will move towards a standard of integrated SNS and DNS analysis. To enable this, a comprehensive catalogue of DNS signatures similar to that of SNS signatures (Wellcome Trust Sanger Institute 2016) would be required.

## Materials & Methods

### Cell line exposure and whole-genome sequencing

MCF-10A and HepG2 cells were obtained from the ATCC. MCF-10A was cultured in DMEM/F12 medium supplemented with 10%,FBS, 10 ng/mL insulin, 20 ng/mL EGF, 0.5 μg/mL hydrocortisone, 50 ng/μL penicillin and 50 U/mL streptomycin. HepG2 was cultured in minimal essential medium (MEM) supplemented with 10%,FBS, nonessential amino acids, 50 ng/μL penicillin and 50 U/mL streptomycin. For cisplatin exposure, 60,000 (MCF-10A) or 250,000 (HepG2) cells/well were seeded at day 0 in a 6-well plate. On day 1 cisplatin was added to final concentrations of 0.5 μM and 1 μM (MCF-10A) or 0.75 μM (HepG2). At day 7, cells were trypsinized, counted, and re-seeded in a new 6-well plate. This process was repeated 8 times. As mutagenesis requires DNA replication, the proliferation rate was monitored (Supplemental Fig. S31). After 4 and 8 weeks, cells were expanded, and single cells were FACS-isolated directly into a 96-well plate with culture medium. These single cell clones were expanded for DNA isolation and whole-genome sequencing. In addition, the MCF-10A and HepG2 cell lines were sampled at the start of the cisplatin exposure. DNA isolation was performed using the Wizard Genomic DNA Purification Kit (Promega, Madison, WI, USA) according to the manufacturer’s instructions. Paired end sequencing was performed on a HiSeq 10x instrument with 150bp reads at Novogene Co., Ltd. (Beijing, China).

### Alignment and variant calling

Read alignment to hs37d5 was done using BWA-MEM, followed by PCR duplicate removal and merging using Sambamba (v0.5.8) (Tarasov et al. 2015). Variant calling was performed using Strelka (v1.014) (Saunders et al. 2012). Variants in dbSNPv132, 1000 genomes (1000 Genomes Project Consortium 2015), segmental duplications, microsatellites and homopolymers, and the GL and decoy sequences were excluded. Additionally, variants were filtered for having at least: 20% variant allele frequency, 25x coverage in both treated and control sample and at least 4 reads supporting the variant. 0.4% and 0.2% of the variants were shared between the clones from the 0.5 μM and 1 μM treated cells. Supplemental Fig. S32 shows the variant allele frequency distribution.

DNSs were identified as 2 adjacent SNSs. As primary QC we checked that the variant allele frequencies of both SNSs were equal. Secondly, we re-called the genomes using Freebayes, which calls DNSs when the SNSs are in the same reads (Garrison 2012). Out of the 2,868 DNSs extracted from the Strelka calls, 2,818 were also called by Freebayes. Lastly, we checked the DNSs in IGV. All DNSs identified from the Strelka analysis were in the same DNA molecule. Focusing specifically on those DNSs that were not called by Freebayes, 17 were not called as DNSs by freebayes as they were close to a germline variant, and Freebayes called these as tri- or tetranucleotide substitutions. Beyond this, 24 putative DNSs were part of complex mutations that were not in fact DNSs. Of the remaining 9 putative DNSs not called by Freebayes, 5 were likely false-positives, as most were only present in one sequence read direction, in regions with low mapping quality, or located near the end of sequencing reads. Overall, we estimated the initial false-discovery rate of DNSs to be ~1.2% (33/2,868) but after Freebayes and IGV inspection we estimate that the false-discovery rate is close to zero. For indels the intersection between the Strelka and Freebayes calls was used.

### Copy number analysis

Freebayes calls were filtered to select variants in dbSNPv132. Coverage and B-allele-frequencies were extracted and segmented using the Quantsmooth (v1.44.0) package in R (Eilers and de Menezes 2005).

### Detection of Structural Variants

Manta v0.29.6 was used to detect structural variants (SVs) present in the cisplatin-treated but not the untreated samples (Chen et al. 2016). The following filters were applied: 1) Breakpoints of intra-chromosomal SVs must be >= 1000 base pairs apart. 2) Both breakpoints must be located on autosomes. 3) Each candidate SV must be supported by at least 10 spanning or split reads.

### Statistical analysis of enrichment of extended sequence context

To test for enrichment or depletion of SNSs in extended sequence context we used a binomial test. The null hypothesis was that the proportion of occurrences of a given penta- or heptanucleotide centered on a given SNS (one of C>A, C>G, C>T, T>A, T>C or T>G) was the same as the proportion of ***all*** penta/heptanucleotides centered on that SNS.

We take as an example a C>T SNS at the center of pentanucleotide TC<u>C</u>AT in the combined MCF-10A data. There were a total of 9,509 C>T mutations in the sequenced portions of the genome, of which 162 were in TC<u>C</u>AT. In total there were 1,089,134,720 pentanucleotide sites centered on C in the sequenced regions of the genome, of which 7,046,748 were TC<u>C</u>AT. We then used the R function call

binom.test(x = 162, n=9,509, p = (7,046,748 / 1,089,134,720)),

which rejected the null hypothesis with *p* < 1.13×10^-26^.

### Analysis of association between cisplatin mutations and genomic features

We obtained processed ChIP-seq datasets for HepG2 for H3K4me1, H3K4me2, H3K4me3, H3K9ac, H3K9me3, H3K27ac, H3K27me3, H3K36me3, H3K79me3, CTCF and EZH2 from www.ncbi.nlm.nih.gov/geo/, accession GSE29611. As histone ChIP-seq data for MCF-10A was not available, we obtained analogous data for normal human mammary epithelial cells and used this a substitute. We obtained MCF-10A expression data from (www.ncbi.nlm.nih.gov/geo/, accession GSM1100206), and HepG2 expression data from the Epigenomic Roadmap (http://egg2.wustl.edu/roadmap/data/byDataType/rna/expression/57epigenomes.RPKM.pc.gz). For analysis of replication timing and replication strand bias, we obtained processed replication timing (RepliSeq) data for HepG2 and MCF7 (MCF-10A data was not available) from GEO, (www.ncbi.nlm.nih.gov/geo/, accessions GSM923446 and GSM923442). We determined replication strand according to (Liu et al. 2016).

### Sources of publicly available sequencing data

This study used whole genome sequencing data from 264 HCCs from Japan (Fujimoto et al. 2016) and 78 from Hong Kong (Kan et al. 2013) and 140 ESADs (Noorani et al. 2017). Additionally, we used whole genome sequencing data of 24 lung adenocarcinomas (Imielinski et al. 2012) and 112 melanomas. For the HCCs, ESADs and melanomas, simple somatic mutation data was downloaded from the ICGC data portal (https://dcc.icgc.org/, release 18, March, 2015). The 78 Hong Kong HCCs were re-analyzed as described previously (Huang et al. 2017).

### Analysis of the SNS cisplatin signature exposure in tumors

We used the mSigAct signature presence test (Ng et al. 2017) to assess presence of the experimental cisplatin SNS signature in the publicly available mutational spectra of HCCs and ESADs as specified in Supplemental Data S2. Briefly, the mSigAct signature presence test determines the likelihood of the observed mutation spectrum with and without a contribution from the target signature and compares these with a likelihood ratio test. The null hypothesis is that the counts are generated without a contribution from the target signature and the alternative hypothesis is that they were generated with a contribution from the target signature. We took the weighted average of the SNS spectra of all MCF-10A and HepG2 cisplatin clones as the cisplatin SNS signature. As described in (Ng et al. 2017), the mSigAct signature presence test has better receiver operating characteristics than the NMF approach from (Alexandrov et al. 2013a; Alexandrov et al. 2013b) (LA-NMF for short), as assessed by tests on simulated data. More concretely, the mSigAct signature presence test is better suited for conservative assessment of the presence of a signature. In what follows, we will use the customary notation for NMF, *V* ≈ *WH,* in which *V* is the matrix of observed mutational spectra, *W,* is the matrix of mutational signatures, and *H* is the matrix of “exposures”. LA-NMF imposes no sparsity constraints on the number of signatures operating in a tumor (i.e. on the number of non-zero elements in columns of *H*). As shown in Section 4.3 of (Alexandrov 2014), the *W* and *H* matrices determined by LA-NMF are sometimes highly variable depending on the specific subset of tumors in *V,* especially when *V* contains relatively small number of tumors, as is the case for the current study. The mSigAct presence test avoids this problem by using the precomputed consensus signatures computed by LA-NMF to address the narrower question of whether a given signatures is plausibly necessary to account for a single tumors mutational spectrum. The mSigAct software is available from the URL https://zenodo.org/record/843773#.WZQQE1EjHRZ as the following doi: 0.5281/zenodo.843773 (Ng et al. 2017).

### NMF on DNS spectra

To assess the effect of cisplatin on primary tumors based on DNSs, we developed a customized semi-supervised NMF (ssNMF) method that incorporated the method from (Schmidt 2007) into the LA-NMF code from (Alexandrov et al. 2013b); Supplemental Data S3 provides the patch file. Again using the notation *V* ≈ *WH* to describe NMF, ssNMF treats *W,* the signature matrix, as composed of two segments: *W_f_,* which specifies the known, fixed signatures, and *W*_*u*_, which is computed by NMF. ssNMF updates only *W_u_* and *H.* The advantage of using ssNMF rather than the closely related method in (Alexandrov et al. 2013b) is that ssNMF can directly ask the question: "To what extent can the DNS spectra of sets of HCCs and ESADs be explained by the action of the experimental cisplatin DNS signature combined with a reasonably small number of additional, unknown signatures?" By contrast, NMF would have to rediscover the cisplatin signature, which, because of the inherent limitations of LA-NMF, may vary from the experimental signature (Alexandrov 2014). Supplemental Data S4 describes in more detail the advantages of using ssNMF.

We ran ssNMF separately on (i) *V*_*HCC*_, which contained the DNS spectra of the HCCs, lung adenocarcinomas, and the cell line clones and (ii) *V*_*ESAD*_, which contained the DNS spectra of the ESADs and the cell line clones. We ran ssNMF on each of *V_HCC_* and *V*_*ESAD*_, asking for 2, 3, 4 and 5 signatures in total. In all cases *W_f_* consisted of a fixed signature that was the weighted average of the DNS signatures of MCF-10A and HepG2. Using the signature stability and average Frobenius reconstruction error approach described in (Alexandrov et al. 2013b), we chose 3 signatures for both *V_HCC_* and *V_ESAD_* (Supplemental Fig. S25, S26). As a further sanity check on the ssNMF analyses, we ran analogous analyses using LA-NMF. This yielded very similar results (Supplemental Data S4).

## Data availability

Sequencing reads for the cisplatin exposed MCF-10A and HepG2 clones are available at the European Nucleotide Archive (http://www.ebi.ac.uk/ena) under accession number PRJEB21971. The sample accessions are listed in Supplemental Table S1.

## Acknowledgements

We thank Willie Yu for technical assistance.

This study was funded by NMRC/CIRG/1422/2015 to SGR.

## Author contributions

AB and SGR designed the study. AB and SZH performed *in vitro* experiments. AB, MNH and JQL carried out electronic analyses. AWTN provided analysis tools and technical support. YK, KC and HN provided clinical information. AB and SGR drafted the manuscript and prepared figures. AB, BTT and SGR edited the manuscript. All authors read and approved the final manuscript.

